# Genetic consequences of effective and suboptimal dosing with mutagenic drugs in a hamster model of SARS-CoV-2 infection

**DOI:** 10.1101/2023.02.20.529243

**Authors:** Christopher J. R. Illingworth, Jose A. Guerra-Assuncao, Samuel Gregg, Oscar Charles, Juanita Pang, Sunando Roy, Rana Abdelnabi, Johan Neyts, Judith Breuer

## Abstract

Mutagenic antiviral drugs have shown promising results against multiple viruses, yet concerns have been raised about whether their use might promote the emergence of new and harmful viral variants. Here, we examine the genetic consequences of effective and suboptimal dosing of favipiravir and molnupiravir in the treatment of SARS-CoV-2 infection in a hamster model. We identify a dose-dependent effect upon the mutational load in a viral population, with molnupiravir having a greater potency than favipiravir per mg/kg of treatment. The emergence of de novo variants was largely driven by stochastic processes, with evidence of compensatory adaptation but not of the emergence of drug resistance or novel immune phenotypes. Effective doses for favipiravir and molunpiravir correspond to similar levels of mutational load. Combining both drugs had an increased impact on both efficacy and mutational load. Our results suggest the potential for mutational load to provide a marker for clinical efficacy.

## Introduction

Molnupiravir (EIDD-2801/MK-4482) is an oral prodrug of the nucleoside analogue β-D-N4-hydroxycytidine (EIDD-1931, NHC) with broad inhibitory activity against RNA dependent RNA polymerases of positive and negative strand RNA viruses (RdRps) including SARS-CoV-2 and other coronaviruses^1–6^. Following oral administration, molnupiravir is metabolized and phosphorylated into the active drug NHC 5’-triphosphate (NHC-TP). During viral replication NHC-TP is substituted by SARS-CoV-2 RdRp for cytidine or uridine triphosphate into newly synthesized viral RNA, resulting in incorporation of either guanosine or adenosine when the RNA is copied^3,7^. The accumulation of random C to U or G to A transition errors in the viral RNA genome is associated with lethal mutagenesis resulting in loss of viral fitness as evidenced by reduced viral loads, reduced viral infectivity and reduced lung pathology in animal models^8,9^. Favipiravir, a nucleoside analogue, has also been associated with C to U and G to A transition errors and lethal mutagenesis^8,10^. However a number of small clinical trials have not shown that favipiravir is able to reduce SARS-CoV-2 viral loads or prevent disease progression^11^.

When administered in a randomised phase 3 clinical trial to SARS-CoV-2 positive individuals who were symptomatic for five days or fewer, molnupiravir taken for five days was reported to reduce rates of hospitalisation and death^12^. Against a background of concerns about efficacy, it has been licensed by the Food and Drug Administration (FDA) for use in mild to moderate coronavirus disease (COVID-19) where the person was at increased risk for progression to severe disease, and under similar conditions for use in the UK^13–15^. A recent clinical trial has shown that molnupiravir increased the transition:transversion mutation ratio in SARS-CoV-2^16^. The potential use of molnupiravir in combination with favipiravir has also been suggested^17^.

Concerns have been raised that the mutagenic mode of action of molnupiravir could drive the emergence of drug-resistance mutations against itself or other antiviral therapies^18,19^. Cases of the emergence of resistance to antiviral drugs have been reported where treatment has failed to clear SARS-CoV-2 infection^20,21^. Understanding viral behavior given suboptimal dosing of mutagenic drugs is a research priority.

We previously used a Syrian hamster model to demonstrate the efficacy of short term (4 days) molnupiravir treatment on reduction of SARS-CoV-2 viral load, infectivity and lung pathology^22^. In this model, a dose of 200 mg/kg BID was effective in reducing infectious virus in the lung by 1.8 to 2.0 log_10_ TCID_50_/mg lung tissue (depending on the SARS-CoV-2 variant used for challenge), with evidence of significantly reduced lung pathology in treated animals. In similar experiments, treatment with 600 mg/kg BID of favipiravir for four consecutive days achieved a reduction of nearly 2 log_10_ in infectious virus titers in the lungs. Here, using viral sequence data from these hamster studies, we examine the impact of effective and suboptimal dosing with molnupiravir (200mg/kg BID and <200 mg/kg BID respectively)^23^ and favipiravir (600 mg/kg BID and <600 mg/kg BID respectively)^24^ upon genetic diversity in the viral populations. We assess the likely consequences of the transmission of variants induced by mutagenic treatment, and examine the potential emergence of drug resistance under treatment. By comparing data from animals treated with different mutagenic drugs, we consider the potential for sequence-based statistics to be used as a biomarker for clinical efficacy.

## Results

Both molnupiravir and favipiravir target viral replication, leading to an increase in the effective mutation rate of the virus and consequently in mutational load. Mutational load was calculated in terms of the sum of minor allele frequencies across sites in the genome; increases in the mutation rate of the virus are expected to increase this statistic under a model of mutation-selection balance^25^. Linear regression models fitted to our data showed significant increases in the extent of mutational load in samples from hamsters treated with increasing doses of drug for both favipiravir and molnupiravir (p-values < 10^−7^) (Figure 1A,B).

**Figure 1:**
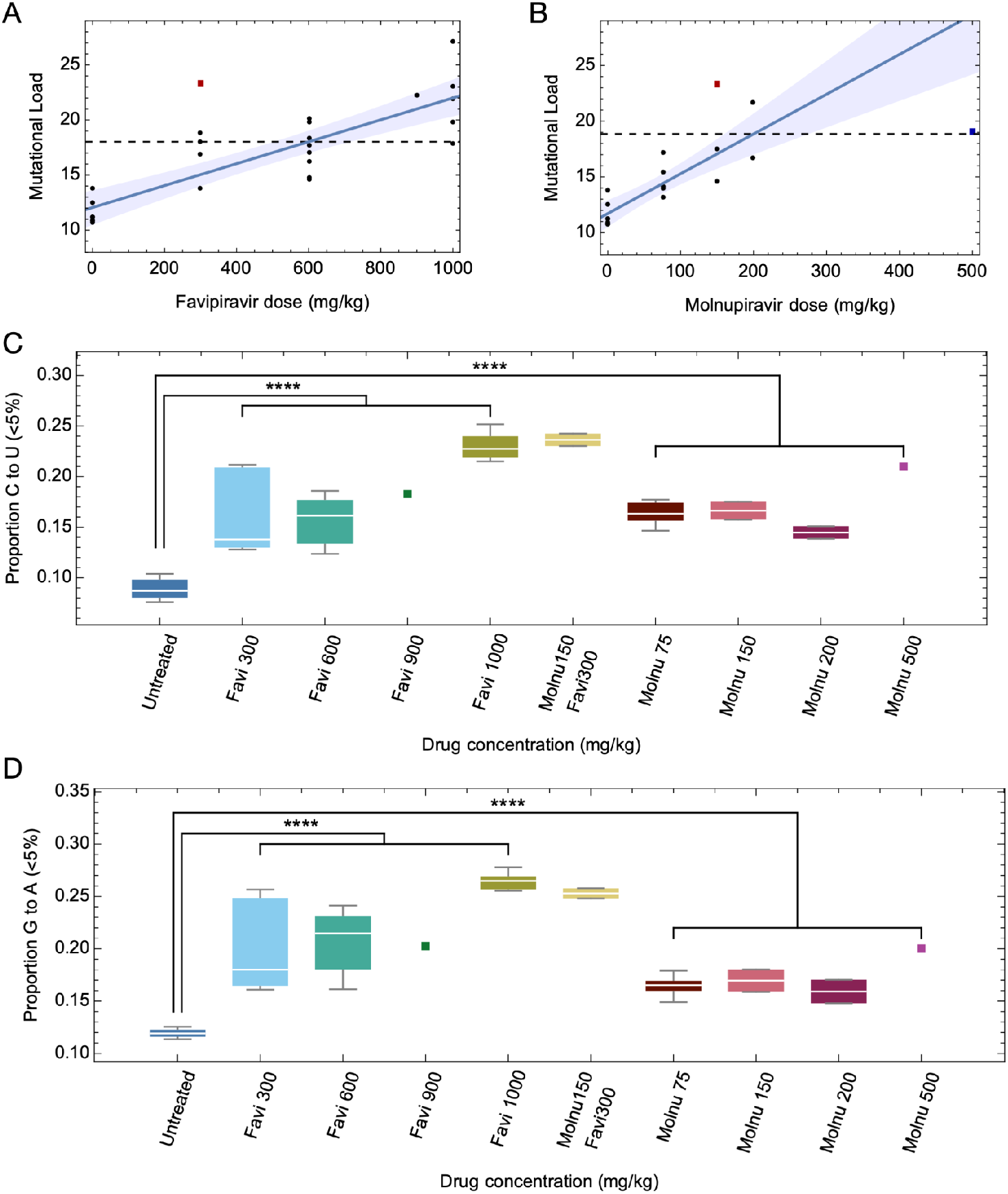
Statistics of mutational load in treated and untreated populations. **A**. Increases in mutational load were observed in viral populations treated with increasing doses of favipiravir. Black dots show genome-wide statistics calculated from populations. The red square shows the mean value for populations treated with 300mg favipiravir and 150mg molnupiravir. The blue square at 500mg molnupiravir shows a datapoint that was omitted from the regression calculation; this point is discussed in the main text. Shading indicates a 95% confidence interval for the regression line. A horizontal dashed line indicates the model threshold at the point of effective treatment. **B**. Increases in mutational load were observed in viral populations treated with increasing doses of molnupiravir. **C**. The proportion of low frequency variation (variant frequency <5%) that was comprised of C to U mutations was higher for treated populations than for untreated populations. Squares show treatments for which a single data point was collected. **D**. The proportion of low frequency variation that was comprised of G to A mutations was higher for treated populations than for untreated populations

The use of both favipiravir and molnupiravir have previously been associated with decreases in viral infectivity^22,24^. According to our analysis, molnupiravir is more potent than favipiravir in this respect, with a greater extent of mutational load induced per mg of drug (p=9 × 10^−5^, Students t-test). Samples collected following combination of 150mg/kg BID molnupiravir and 300mg/kg BID favipiravir treatment had a greater mutational load than samples collected following the use of either of the constituent parts of the combined treatment (p-values 0.02 and 0.04 respectively, student’s T-test) (Figure 1A,B). This increase in mutational load suggests that the two drugs operate in a synergistic way. Where either drug, given singularly, caused an increase in the viral mutation rate, we suggest that the combination of two drugs is likely to have resulted in a significant additional increase in the rate of mutation. An analysis of sequence data found no correlation between read depth and mutational load, suggesting that our results were not the result of artefacts of the experimental process (Supplementary Figure 1).

When evaluated in terms of mutational load, the thresholds at which drug dosing became effective for SARS-CoV-2 treatment in the hamsters (i.e. 600 BID mg/day and 200 BID mg/day for molnupiravir)^22,24^ were very similar, with 18.0 mutations per genome for favipiravir, and 18.9 mutations per genome for molnupiravir in our regression model (Figure 1A,B). Given the uncertainty in our model, and the imprecise measurement of thresholds for efficacy, these two values are statistically inseparable. Evolutionary theory suggests that viral fitness is directly related to the amount of mutational load^25^, highlighting this statistic as being of potential utility as a clinical marker to assess the efficacy of mutagenic antiviral drugs.

Our linear regression analysis excluded one data point, collected from a population treated with 500mg/day of molnupiravir. Under a simple model, the extent of mutational load achieved by a population will be proportional to the mutation rate of the virus^25^. However, achieving a high mutational load requires a population to remain alive for sufficiently long to generate viruses with that quantity of mutations. In the excluded case, we propose that mutagenesis caused the extinction of the population substantially before it reached mutational load equilibrium. The collection of only a single data point at this dose prevented further investigation of this phenomenon.

Both favipiravir and molnupiravir caused a significant change in the mutational spectrum of the SARS-CoV-2 virus (Figure 1C,D). A previous study examining the effect of favipiravir during influenza B used the composition of variants with frequencies less than 5% as a marker for the effect of a mutagenic drug^10^. Applying the same statistic, we identified significant increases in the proportions of C to T and G to A mutations among samples from hamsters treated with either drug versus samples from untreated animals (p<1×10^−6^, Student’s t-test) (Figure 1C,D).

Together these results suggest that favipiravir and molnupiravir cause viral mutagenesis even at suboptimal doses, with treatment using a higher dose being distinguished by a greater mutagenic effect on the viral population. When given together, the combined therapy leads to an effect greater than that of either of the constituent drugs at the same dose. Our findings can be understood in terms of theoretical predictions about the behaviour of mutagenic drugs^26^. The viral mutation rate, and hence the extent of mutational load, increase with increasing concentration of the drug. To substantially reduce viral load the drug has to generate enough mutations to overcome the intrinsic fecundity of the virus. However, where the drug dose is not sufficient to reduce viral load, an effect on the composition of the viral population will still be seen.

We next examined the viral populations for the emergence of variants that could confer phenotypic benefits to the virus, such as resistance to drug therapy. Considering variants that were observed at frequencies of at least 5%, that occurred within open reading frames of the virus, and were supported by a read depth of at least 100, we identified a total of 306 *de novo* variants in the viral genome within the treated populations (Figure 2). Of these variants 207 were non-synonymous with 98 synonymous and one nonsense mutation. Accounting for the proportion of sites that could engender synonymous or non-synonymous mutations, these values indicate a bias towards synonymous variation, with a πN/πS ratio of 0.63 suggesting purifying selection against non-synonymous mutations. An equivalent calculation for variants in untreated populations gave a lower πN/πS ratio of 0.25 (p<0.02, likelihood ratio test, Figure 3), suggesting an increased level of selection against non-synonymous variants in untreated populations. This result is consistent with a signal of compensatory viral adaptation under mutagenic treatment. Theoretical and experimental research in evolutionary biology suggest that reductions in the fitness of a population confer an increased potential for the gain of beneficial variants^27,28^. In this case, as antiviral therapy induces large numbers of deleterious mutations, disrupting viral protein function, the number of potential non-synonymous variants which restore viral fitness is increased. Non-synonymous mutations that were strongly selected against in a fit viral population become, under therapy, less bad for the virus.

**Figure 2:**
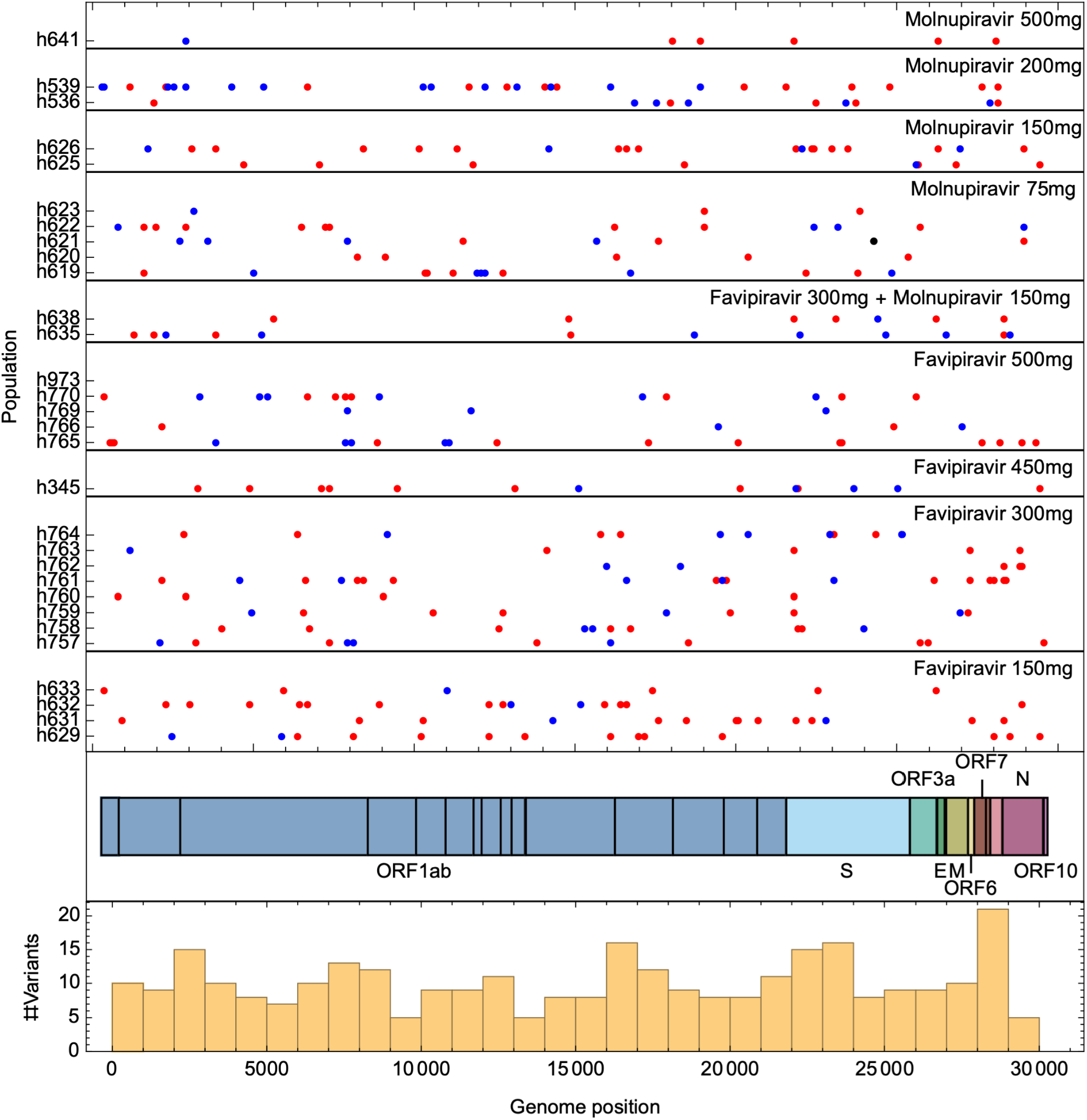
Genomic locations of variants in treated viral populations. The locations of variants which reached a frequency of at least 5% in the viral population are shown in red (non-synonymous variant), blue (synonymous variant) or black (nonsense variant). With the exception of variants transmitted through standing variation, very little replication of variants between treated populations was observed.

**Figure 3:**
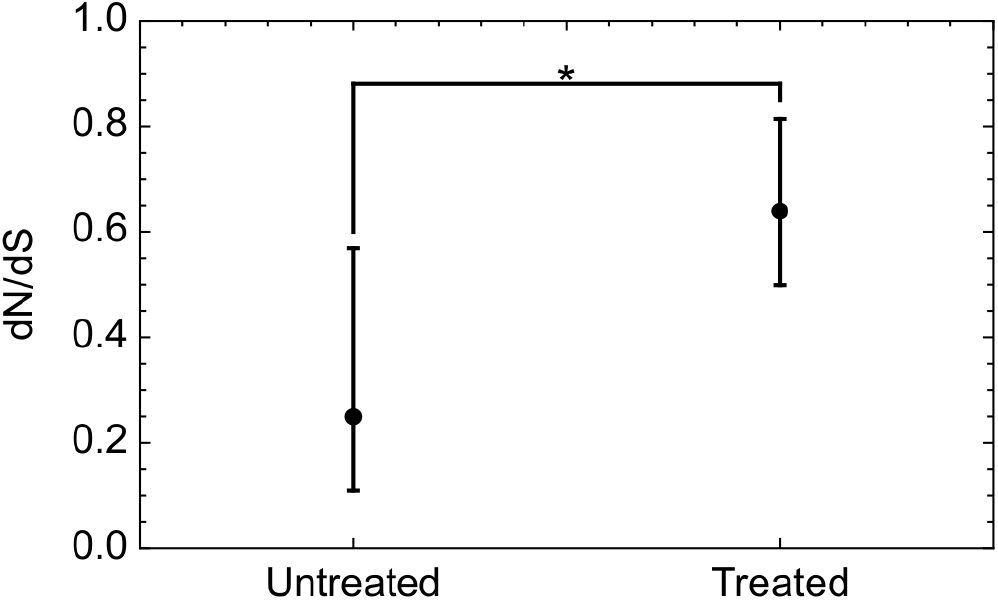
dN/dS was lower in untreated than in treated populations. Error bars were calculated using a likelihood model.

The potential signal of compensatory adaptation was not accompanied by evidence of the adaptive gain of mutations conferring specific fitness gain. Of the variants observed in the 30 treated populations, only six were observed in more than one of the populations, suggesting a highly stochastic process of variants emerging to frequencies of 5% or more. As a potential exception to this, the A21804G variant, which codes for the N81S mutation in the Spike protein, was observed as a consensus level substitution in two populations that were treated with favipiravir. *De novo* variants were roughly uniformly distributed across the genome, following a pattern seen in the general evolution of SARS-CoV-2 in the human population^29^, and with no clear bias towards any one part of the genome. Despite our finding of a link between mutational load and drug dose, there was no significant relationship between treatment type and the number of *de novo* variants observed at higher frequencies; the expansion of new variants to detectable frequencies was dominated by stochastic events (Supplementary Figure 2).

We further did not identify evidence of adaptation that could confer resistance against the antiviral drugs used in these experiments. Upon 5’-triphosphorylated of the parent nucleoside of remdesivir, the molecule is incorporated by the RdRp into the growing RNA chain and allows for addition of three more nucleotides before RNA synthesis stalls^30^. The variants observed in hamster populations at frequencies of 5% or greater did not replicate known remdesivir-resistance variants observed in SARS-CoV-2^20,31,32^. Two consensus-level variants were observed in RdRp from hamsters given treatment with favipiravir, but no consensus-level variants were observed in RdRp from hamsters given treatment with molnupiravir (Figure 4). No mutations in the RdRp binding site, or in sites that have been associated with drug resistance for other RdRp inhibitors, were found (Supplementary Table 1). Within the Spike protein consensus-level variants were observed in populations treated with favipiravir, two of which coded for the N81S substitution, while a single consensus-level variant was observed for populations treated with molnupiravir (Supplementary Figure 3, Supplementary Table 2). One variant, I692V, observed at a frequency of 8% in a population treated with molnupiravir, was among the set of variants which defined the B.1.1.198 variant of concern, identified early in the pandemic^33^.

**Figure 4:**
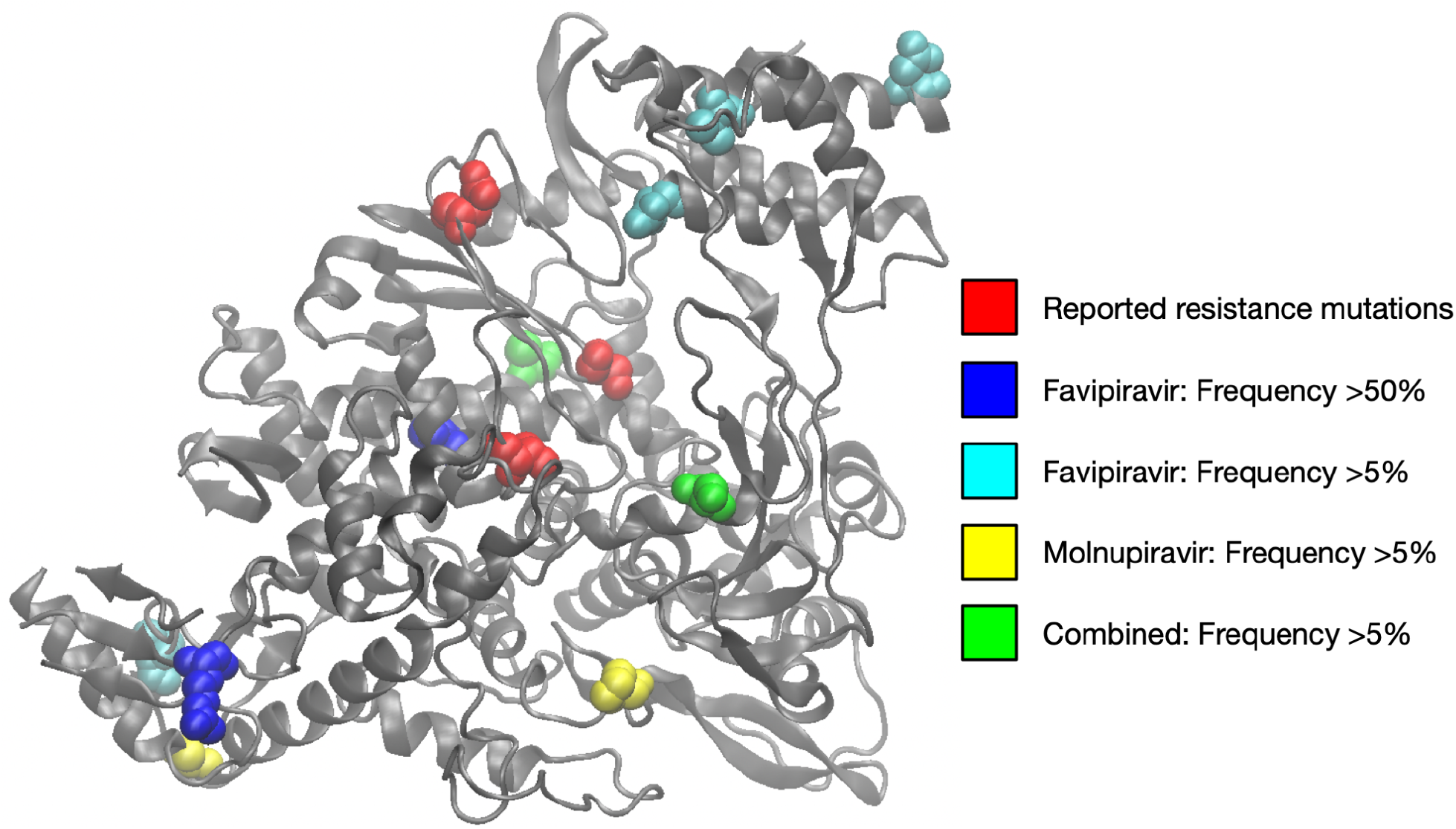
Variants observed in RdRp at frequencies of 5% or greater. Sites reported to convey resistance to remdesivir are shown in red vdW representation^20,31,32^. Other colours show variants observed in hamster populations at frequencies of 5% or more. Image created in VMD^42^ based on the PDB structure 6m71^43^

## Discussion

Both molnupiravir and favipiravir are mutagenic antiviral drugs. When used against SARS-CoV-2 they interfere with viral replication, leading to an increase in the mutation rate of the virus. Increasing the mutation rate raises the possibility that consequent increased genetic variation will result in the evolution of drug resistance mutations, or of mutations that could make a virus better adapted to a new host. Previously, a SARS-CoV-2 infection model in Syrian hamsters showed a dose-dependent reduction in viral loads, infectivity and lung pathology in molnupiravir or favipiravir treated animals^22–24^. Taking this further, we here used viral genome sequence data to explore how both favipiravir and molnupiravir shaped the viral populations in these animals.

Our identification of a pattern of increase in mutational load with increasing drug dose highlights the fact that even suboptimal doses of each drug are likely to affect the viral population. Given a sufficient dose, treatment induces mutations to the point that viral load is significantly reduced^22,24^. However, even suboptimal treatment increases mutational load. Irrespective of the level of treatment, viruses transmitted from a treated host are therefore likely to carry an increased number of mutations. The consequences of this increased number of mutations would depend upon the specific mutations the virus acquired. In a situation where all mutations were deleterious, any increase in the number of viral mutations would be of potential clinical benefit; a virus carrying deleterious mutations could potentially cause a milder case of infection. Recent work has shown a potential effect of mutagenic drugs in SARS-CoV-2 populations, and an accumulation of mutations in patients treated with antiviral drugs^34,35^. We note that the acquisition of mutations by the virus is not intrinsically problematic for human health; negative consequences arise only when such mutations drive increased pathogenicity or virulence.

The dose-dependent response to therapy we identify is consistent with a pattern in which accumulated mutations generally reduced viral fitness. This conclusion is supported by the finding of selection against non-synonymous mutations, and is consistent with the dose-dependent reduction in viral infectivity and lung pathology observed experimentally for both favipiravir and molnupiravir^23,24^. In combined molnupiravir and favipiravir treatment, a significant reduction in viral infectivity^17^ was mirrored by a significant increase in mutational burden. Where mutated viral genomes survive, they likely do so despite a fitness that is lower than the pre-treatment wild type.

Our study identified a reduced bias against non-synonymous variation in treated as opposed to untreated populations. A lower mean fitness cost of nonsynonymous variants is an expected consequence of reducing the fitness of the virus population, as lower fitness populations have by nature more evolutionary routes via which fitness can be restored^27,28^. This finding provides further indirect evidence for the impact of drug-associated mutagenesis on loss of viral fitness.

The observed lack of repeated variants in treated populations is consistent with a process whereby variants rose to high frequency by chance, via a random process of mutation and genetic drift. Against this background, we cannot exclude the possibility of having observed some genuinely beneficial mutations, whether conferring adaptation to the hamster host or some form of drug resistance. Adaptive viral evolution in multiple directions has previously been observed in experimental cases of animal infection^36^. Thus, a process in which distinct beneficial mutations emerged in different hosts cannot entirely be ruled out.

Our linear regression model suggested that an effective dose of treatment for both favipiravir or molnupiravir corresponded with a mutational load of close to 18 mutations per virus. In so far as mutational load shares a straightforward relationship with viral fitness, this statistic has some promise as a clinical biomarker for evaluating the effectiveness of mutagenic drugs. However, several steps would be required to fully validate its use. Firstly, the values measured for mutational load in this study incorporate an error profile that is specific to the sequencing process used. Secondly, in the data collected, identical doses of drug led to variable levels of viral load; further investigation of the extent of this variation is required. Thirdly, hamsters likely represent a more homogeneous population than hospitalised patients. Intrinsic viral fecundity in the absence of treatment may vary according to the particular immune response of an individual, as well as the genotype of the virus.

We acknowledge some limitations to our study. For example, the collection of viral genome sequence data was only possible under conditions that allowed the viral population to be recovered at levels sufficient for sequencing to be performed. A bias could exist towards populations that experienced unusually low levels of viral mutagenesis for a given level of treatment. We imposed a read depth threshold to the data, and examined only samples for which high quality sequence data was obtained. Secondly, data were collected at only one point in time, four days after treatment with antiviral drugs. To this extent we were not able to assess whether the levels of mutational load observed were close to or far from equilibrium, or what was the pattern of change in mutational load over those four days. The manner via which mutations accumulate during treatment is of interest in considering the consequences of virus transmission from a treated patient: A more rapid process of mutation accumulation is likely to increase the mutational burden of any viruses transmitted. Mutation accumulation has been studied in more detail in a case of the extended treatment of an immunocompromised child with RSV infection; in that case, the acquisition of mutational load was rapid, with 60% of the increase in mutational load occurring within the first four days of treatment^37^. The serial collection of viral samples would provide better insight into these processes.

Despite some limitations, our results suggest that treatment with mutagenic drugs increased the extent of mutational load in our viral populations in a roughly continuous manner, with the increase in load being proportional to the level of antiviral dosing. To the extent that mutagenic drugs result in a loss of viral infectivity, it can be concluded that the burden of mutations they impose upon the virus is deleterious, with successive gains of mutations being to the detriment of the viral population. On this basis we would expect that, while suboptimal dosing could lead to the evolution of more mutated viruses, the emerged viruses would be mostly of reduced fitness compared to those transmitted from an untreated population. Our study did not find evidence for either the emergence of antiviral resistance, or the systematic emergence of beneficial variants that would militate against the clinical use of mutagenic drugs. In the absence of specific evidence of deleterious effects, clinical studies describing the efficacy (or otherwise) of mutagenic treatments should be given priority in determining the use of antiviral treatments.

## Methods

### Virus samples

The data for this study was generated from a Syrian hamster model designed to demonstrate the efficacy of short term (4 days) molnupiravir and favipiravir in reducing SARS-CoV-2 viral load, infectivity and lung pathology. Details of these experiments have been described in previous publications^22–24^. Sequence data is available from the Sequence Read Archive with accession number PRJNA935666.

### Sequence Analysis

Fastq files were processed with Trimmomatic^38^ using default settings, retaining reads for which both reads in a pair passed filtering. The remaining reads were aligned to the MN908947.3 reference sequence using bwa^39^, before using samtools^40^ to remove reads that did not align to the primary alignment. Further processing was conducted using the SAMFIRE package^41^. Filtering was conducted to remove low quality base pairs (PHRED score <30), following which variants were identified. Data from sites with a read depth of less than 100x was removed from consideration. Variants at greater than or equal to 5% frequency were identified. Samples were kept for analysis if the mean coverage of the viral genome was at least 150x. A full list of samples is provided in Supplementary Table 3.

### Mutational load

Measurements of mutational load were calculated using the aligned sequence data. Under mutation-selection balance, an allele tends to an equilibrium frequency of U/s, where U is the mutation rate and s is the magnitude of selection against the allele^25^; changes in mutation rate thus increase the extent of variation in a genome. We assessed mutational load by calculating the sum of allele frequencies across the genome. At position i in the genome we calculated the frequency q_ia_ of the allele a for each non-consensus allele a, independent of read depth. The mutational load was then calculated as

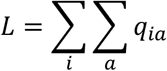

where the sums are taken over all positions in the genome and all non-consensus alleles.

Calculations of mutational load considered all sites in the genome irrespective or read depth, but conditional on the mean read depth of the population. In order to evaluate the effect of sites in the genome with low coverage, we recalculated mutational load values to include only sites with a read depth of at least 50 or at least 100 (Supplementary Figure 4). Linear regression showed that the filtered mutational load values were on average between 1.6% (depth ≥ 50) and 4% (depth ≥ 100) lower than the unfiltered values, with extremely strong linear relationships between the filtered and unfiltered data.

### Analysis of variant composition

Variants were calculated from the data generated in a manner described in the sequence analysis section above. To analyse the composition of variants we calculated πN/πS, defined as

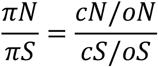

Where cN and cS were the counts of non-synonymous and synonymous variants reaching a frequency of 5% or more in the experimental populations, and oN and oS are the number of potential non-synonymous and synonymous variants that could arise given the various reading frames in the SARS-CoV-2 genome, using the consensus sequence of the initial viral population as a reference. A value of this statistic less than one suggests that the emergence of variants to a frequency of 5% or greater was affected by purifying selection.

To calculate a p-value for the difference between πN/πS values, we calculated likelihoods for the underlying proportions p_U_ and p_T_ of variants in the untreated and treated populations which were non-synonymous, relative to the opportunity, and based upon the observations made. Where we observed cN_U_ nonsynonymous variants and cS_U_ synonymous variants in the untreated population, and cN_T_ nonsynonymous variants and cS_T_ synonymous variants in the treated population, a likelihood for the values p_U_ and p_T_ is given by

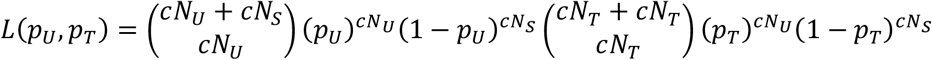

This distribution is shown in Supplementary Figure 5. Defining the hypothesis H_1_: p_U_<p_T_, and H_0_: p_U_≥p_T_, and integrating over the corresponding regions of this distribution, we obtained the p-value given in the main text.

### Identification of spike mutations of interest

As the hamsters were infected with Wuhan-like virus, we assumed mutations of interest would be those identified in the human population in variants of concern / variants of interest. Mutations arising were looked up in tables of mutations with antibody escape information from the following publication and referenced material^33^.

### Visualization of protein structures

Images of protein structures were made with the Visual Molecular Dynamics software package^42^.

## Author contributions

Conceptualisation: CI, JG-A, SG, JB. Data Curation: CI, JG-A, SG. Formal Analysis: CI, JG-A, SG, OC. Funding Acquisition: JB. Investigation: CI, JG-A, SG, OC, JP, SR. Methodology: CI, JG-A, SG, RA. Project Administration: CI, RA, JN, JB. Software: CI, JG-A, SG. Supervision: JG-A, SG, RA, JN, JB. Validation: CI, JG-A, RA, JN. Visualisation: CI, JG-A, SG, OC. Writing - Original Draft Preparation: CI, JG-A, SG, JB. Writing - Review and Editing: CI, JG-A, OC, RA, JN, JB.

