## Supplementary Information for "Genetic consequences of effective and suboptimal dosing with mutagenic drugs in a hamster model of SARS-CoV-2 infection"

### Supplementary Figures

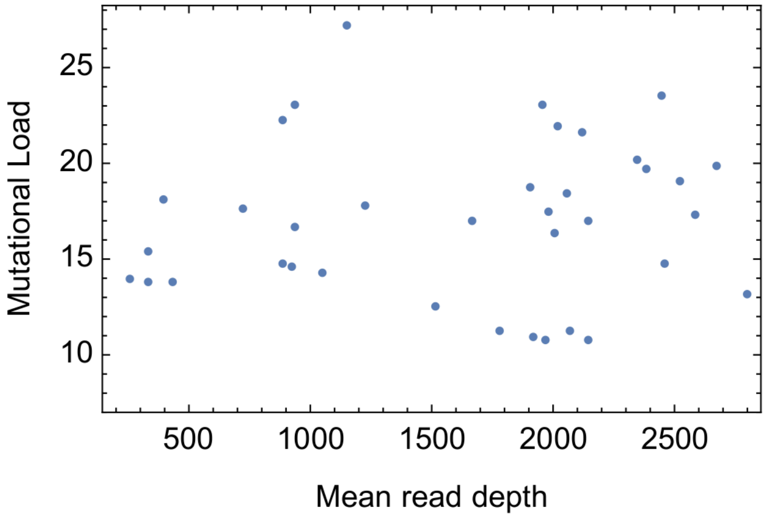


***Supplementary Figure 1:*** *No correlation was observed between the mean read depth of a sample and the extent of mutational load (p-value 0.60, T-test)*

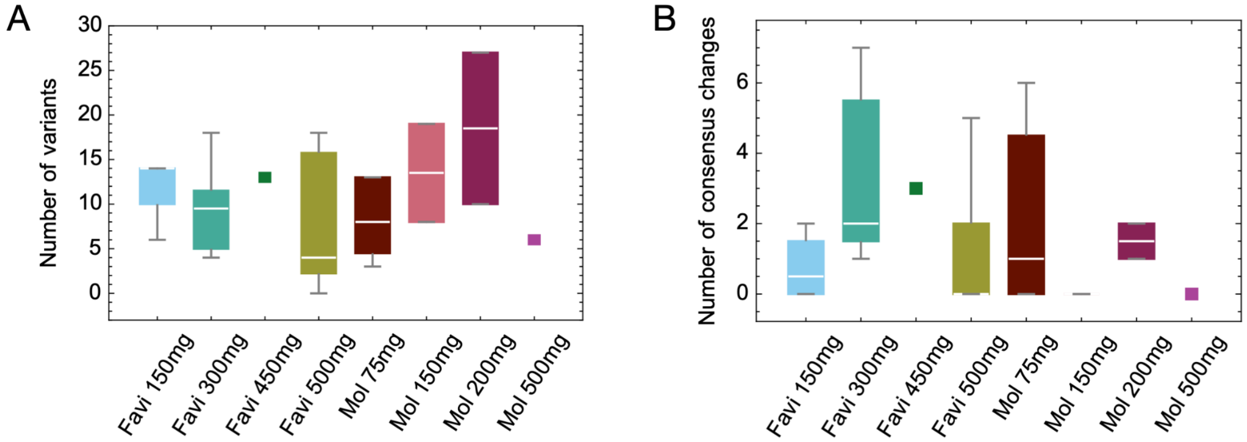


***Supplementary Figure 2: A.*** *Numbers of de novo variants reaching >5% frequency in populations under each treatment.* ***B.*** *Numbers of de novo consensus changes (>50% frequency) in populations under each treatment.*

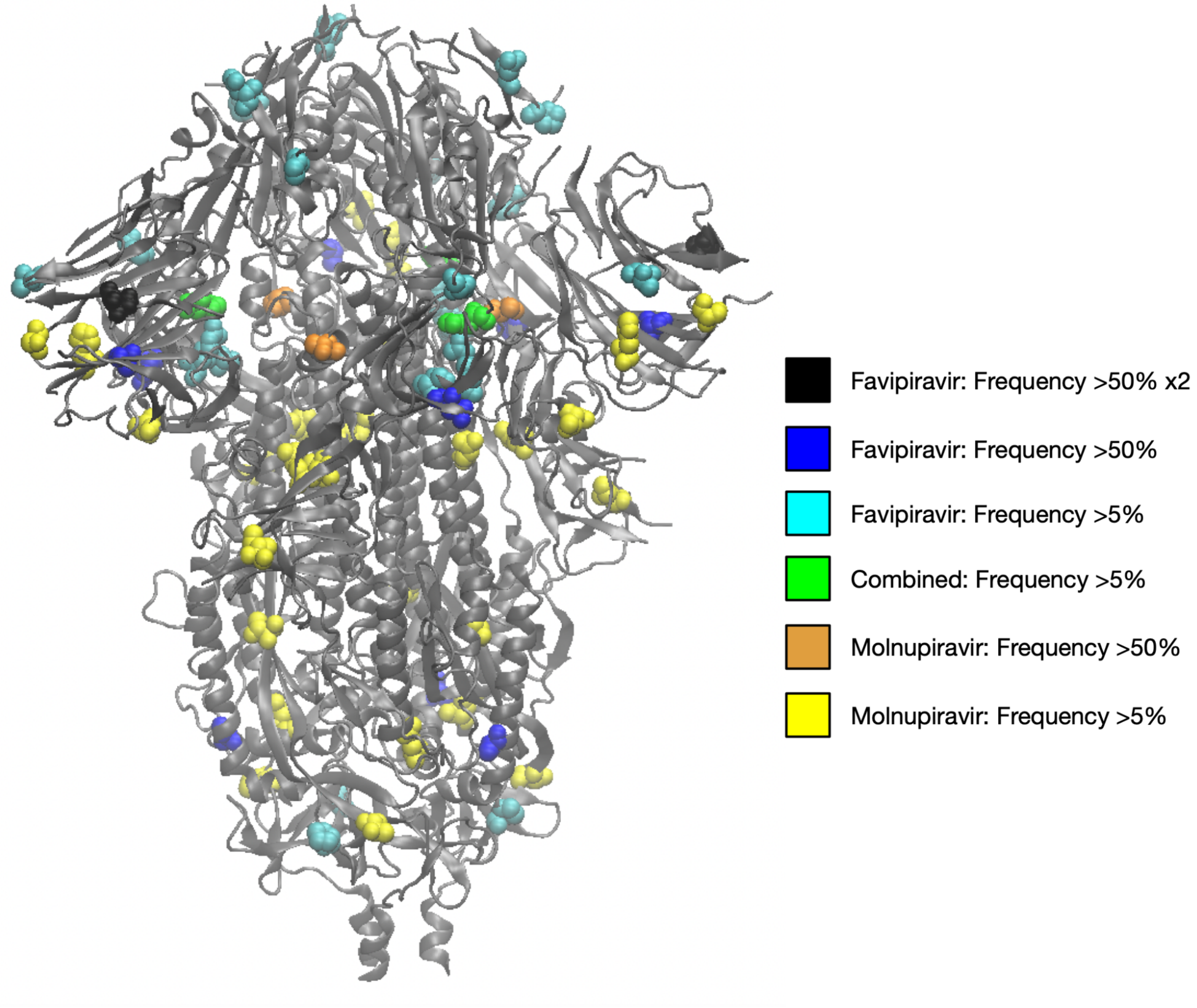


***Supplementary Figure 3: Variants observed in RdRp at frequencies of 5% or greater.*** *Sites reported to convey resistance to remdesivir are shown in red vdW representation[20,31,32]. Other colours show variants observed in hamster populations at frequencies of 5% or more. Image created in VMD[42] based on the PDB structure 6vxx[44]. One nonsynonymous mutation, N81S, was observed in two independent populations treated with favipiravir and is shown in black.*

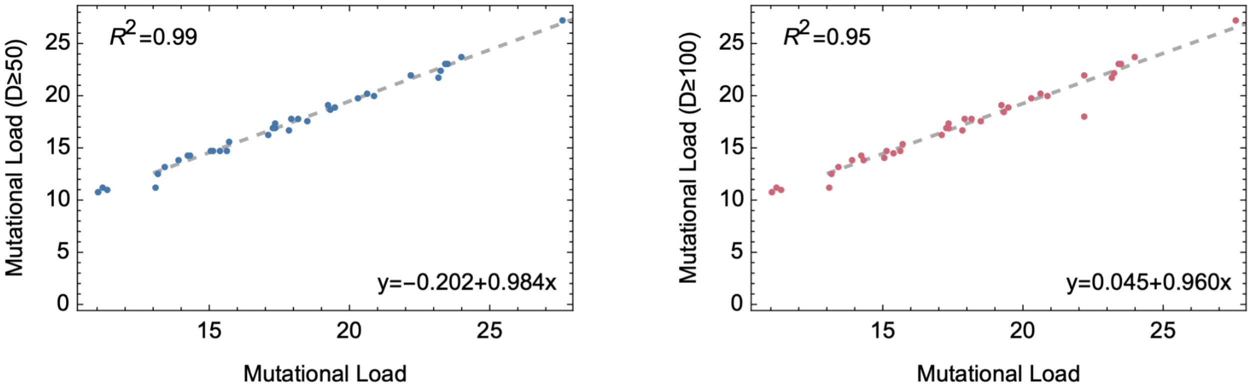


***Supplementary Figure 4: Effect of removing low-coverage sites from calculations of mutational load.*** *Mutational load statistics were calculated removing sites with a read depth of less than 50 (left) or with a read depth of less than 100 (right). The mutational load for removed sites was set to zero, reducing the overall value of the calculated mutational load for each sample. Strong correlations were observed in each case, suggesting that performing the calculation across all genome positions did not led to an overall distortion in the results. Linear regression (shown as a gray dashed line in each case) indicated reductions of close to 1.6% and 4% in mutational load with removal of sites at depths ot less than 50 or less than 100 respectively.*

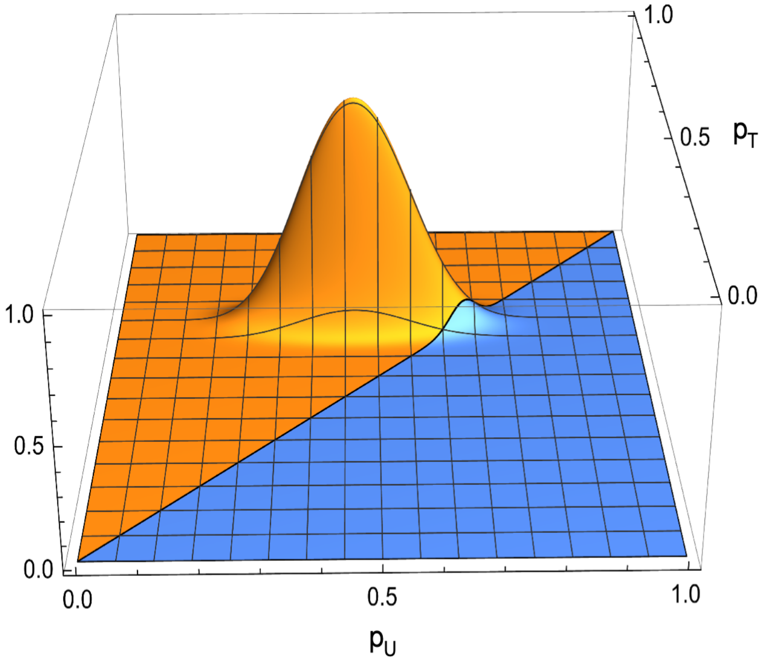


***Supplementary Figure 5: The fraction of non-synonymous variants reaching a frequency of at least 5% was lower in untreated than in treated populations.*** *Joint likelihood function $L(p\text{-}_U, p_T)$ for the underlying proportion of non-synonymous variants in untreated ($p_U$) and treated populations ($p_T$). The relative likelihood, describing the likelihood as a fraction of the maximum likelihood value, is shown. The orange region denotes the section of space for which $p_U<p_T$, while the blue region denotes that for which $p_T \geq p_U$.*
