## Supplementary Table 1 for "Genetic consequences of effective and suboptimal dosing with mutagenic drugs in a hamster model of SARS-CoV-2 infection"

***Supplementary Table 1:*** *Nonsynonymous variants observed in RdRp. Those causing a change in the consensus allele are shown in italic type.*

| ***Nucleotide change*** | ***Amino acid change (RdRp)*** | ***Frequency*** | ***Treatment*** | ***Population*** |
| --- | --- | --- | --- | --- |
| *C13792T* | *R118C* | *0.554* | *Favi 300mg* | *h757* |
| *C14036T* | *A199V* | *0.441* | *Mol 200mg* | *h539* |
| *T14108C* | *I223T* | *0.052* | *Favi 300mg* | *h763* |
| *C14407T* | *P323S* | *0.488* | *Mol 200mg* | *h539* |
| *G14792A* | *S451N* | *0.055* | *Mol 150mg +*  *Favi 300mg* | *h638* |
| *G14885A* | *C482Y* | *0.074* | *Mol 150mg +*  *Favi 300mg* | *h635* |
| *G15793A* | *V785I* | *0.889* | *Favi 300mg* | *h764* |
| *G15937A* | *D833N* | *0.062* | *Favi 150mg* | *h632* |
| *G16078A* | *V880I* | *0.11* | *Favi 300mg* | *h758* |
| *G16117A* | *D893N* | *0.383* | *Favi 150mg* | *h629* |
