## Supplementary Table 2 for "Genetic consequences of effective and suboptimal dosing with mutagenic drugs in a hamster model of SARS-CoV-2 infection"

***Supplementary Table 2:*** *Nonsynonymous variants observed in Spike. Those causing a change in the consensus allele are shown in italic type.*

| ***Nucleotide change*** | ***Amino acid change (Spike)*** | ***Frequency*** | ***Treatment*** | ***Population*** |
| --- | --- | --- | --- | --- |
| *C21575T* | *L5F* | *0.075* | *Mol 200mg* | *h539* |
| *A21788G* | *T76A* | *0.055* | *Mol 150mg +*  *Favi 300mg* | *h638* |
| *A21788G* | *T76A* | *0.076* | *Mol 500mg* | *h641* |
| *A21804G* | *N81S* | *0.979* | *Favi 300mg* | *h760* |
| *A21804G* | *N81S* | *0.786* | *Favi 300mg* | *h763* |
| *C21840T* | *A93V* | *0.593* | *Favi 300mg* | *h759* |
| *C21855T* | *S98F* | *0.051* | *Favi 150mg* | *h631* |
| *C21855T* | *S98F* | *0.191* | *Mol 150mg* | *h626* |
| *G21929A* | *A123T* | *0.139* | *Favi 450mg* | *h345* |
| *C21933T* | *T124I* | *0.054* | *Favi 300mg* | *h758* |
| *C22088T* | *L176F* | *0.71* | *Favi 300mg* | *h758* |
| *C22181T* | *H207Y* | *0.162* | *Mol 75mg* | *h619* |
| *C22338T* | *T259I* | *0.215* | *Mol 150mg* | *h626* |
| *G22346A* | *A262T* | *0.083* | *Favi 150mg* | *h631* |
| *C22419T* | *T286I* | *0.091* | *Mol 150mg* | *h626* |
| *A22503T* | *Q314L* | *0.083* | *Mol 200mg* | *h536* |
| *C22550T* | *P330S* | *0.058* | *Favi 150mg* | *h633* |
| *A22910G* | *N450D* | *0.064* | *Favi 300mg* | *h764* |
| *G23009A* | *V483I* | *0.079* | *Mol 150mg* | *h626* |
| *C23033T* | *P491S* | *0.89* | *Favi 300mg* | *h764* |
| *A23118G* | *H519R* | *0.055* | *Mol 150mg +*  *Favi 300mg* | *h638* |
| *G23262A* | *R567K* | *0.064* | *Favi 500mg* | *h765* |
| *C23277T* | *T572I* | *0.089* | *Favi 500mg* | *h765* |
| *C23277T* | *T572I* | *0.088* | *Favi 500mg* | *h770* |
| *G23318A* | *D586N* | *0.904* | *Favi 500mg* | *h770* |
| *C23453T* | *P631S* | *0.069* | *Mol 150mg* | *h626* |
| *A23636G* | ***I692V*** | *0.082* | *Mol 200mg* | *h539* |
| *C23733T* | *T724I* | *0.242* | *Mol 200mg* | *h536* |
| *C23802T* | *T747I* | *0.886* | *Mol 75mg* | *h619* |
| *C23855T* | *R765C* | *0.063* | *Mol 75mg* | *h623* |
| *C24263T* | *Q901X* | *0.061* | *Mol 75mg* | *h621* |
| *C24333T* | *A924V* | *0.884* | *Favi 300mg* | *h764* |
| *C24789T* | *T1076I* | *0.054* | *Mol 200mg* | *h539* |
| *C24896T* | *P1112S* | *0.072* | *Favi 500mg* | *h766* |
| *G25436A* | *E1262K* | *0.055* | *Mol 75mg* | *h620* |
