## Supplementary Table 3 for "Genetic consequences of effective and suboptimal dosing with mutagenic drugs in a hamster model of SARS-CoV-2 infection"

***Supplementary Table 3:*** *Mean read depths of viral samples*

| ***Population*** | ***Mean Read Depth*** | ***Treatment*** |
| --- | --- | --- |
| h618 | 2068 | Untreated |
| h751 | 1509 | Untreated |
| h752 | 435 | Untreated |
| h753 | 1774 | Untreated |
| h754 | 1968 | Untreated |
| h755 | 2143 | Untreated |
| h756 | 1917 | Untreated |
| h629 | 393 | Favi 150mg |
| h630 | 331 | Favi 150mg |
| h631 | 1903 | Favi 150mg |
| h632 | 2139 | Favi 300mg |
| h633 | 2385 | Favi 300mg |
| h757 | 726 | Favi 300mg |
| h758 | 2342 | Favi 300mg |
| h759 | 925 | Favi 300mg |
| h760 | 1658 | Favi 300mg |
| h761 | 2003 | Favi 300mg |
| h762 | 2056 | Favi 300mg |
| h763 | 880 | Favi 450mg |
| h764 | 889 | Favi 500mg |
| h765 | 939 | Favi 500mg |
| h766 | 2667 | Favi 500mg |
| h769 | 1228 | Favi 500mg |
| h770 | 2021 | Favi 500mg |
| h973 | 1149 | Mol 150mg + Favi 300mg |
| h635 | 2439 | Mol 150mg + Favi 300mg |
| h638 | 1946 | Mol 75mg |
| h619 | 326 | Mol 75mg |
| h620 | 1050 | Mol 75mg |
| h621 | 250 | Mol 75mg |
| h622 | 2582 | Mol 75mg |
| h623 | 2796 | Mol 150mg |
| h625 | 2452 | Mol 150mg |
| h626 | 1982 | Mol 200mg |
| h536 | 939 | Mol 200mg |
| h539 | 2121 | Mol 500mg |
